# *loc*BLAST v2.0 - an improved PHP library for embedding standalone NCBI BLAST+ program to an interactive graphical user interface

**DOI:** 10.1101/556225

**Authors:** T. Ashok Kumar

**Affiliations:** Department of Bioinformatics, Noorul Islam College of Arts and Science, Kumaracoil, Thuckalay 629180, Tamil Nadu, INDIA

**Keywords:** locBLAST, Local BLAST Search, NCBI BLAST+, Sequence Alignment, Embedding BLAST+

## Abstract

**Background:** NCBI BLAST is a most popularly used sequence analysis platform in a broad range of applications. The final release of standalone WWW BLAST server (wwwBLAST v2.2.26) of NCBI was officially released on May 10, 2004, and discontinued its support. Due to the popularity and high demand, the BLAST algorithms were refined with rich features and released as online BLAST, standalone BLAST+, BLAST RESTful service, and cloud BLAST server. *loc*BLAST v2.0 is a lightweight PHP library, designed in preference to WWW BLAST server. It masks command-line BLAST+ programs to a rich graphical user interface (GUI).

**Methods:** The *loc*BLAST v2.0 library was designed using PHP, CSS, pure JavaScript, and standalone NCBI BLAST+ executables.

**Results:** *loc*BLAST v2.0 provides an interactive web interface to input query sequences through the web form and provides a graphical report of the sequence alignment result. The graphical overview, tabular summary, and formatted sequence alignment of *loc*BLAST v2.0 result mimic to the native format of online NCBI BLAST result. It allows users to perform both local and online database search.

**Availability:** Freely available at https://github.com/AshokHub/locBLAST

## 1. Introduction

Sequence alignment is one of the best and well-known method in Bioinformatics for identifying the similarity between the two biological sequences through sequence alignment. Through pairwise sequence alignment, it is possible to predict the structural, functional, and evolutionary relationship of the sequence. The NCBI BLAST (Basic Local Alignment Search Tool) (Altschul et al., 1990) offers different types of programs mainly used for finding local similarity between two biological sequences or a dataset. It provides integrative access to the various biological databases through the web user interface, command-line user interface, and application programming interface. NCBI BLAST+ (Camacho et al., 2009) is a standalone suite of command-line BLAST programs used to compare the query sequence with both online and offline databases. The NCBI BLAST+ executables and source codes (a subset of the NCBI C++ toolkit (Vakatov, 2004)) are freely available at NCBI FTP repository for download.

The standalone NCBI BLAST+ suite consists of various programs and utilities for the different types of sequence alignments, database creation, and data manipulations. The list of NCBI BLAST+ programs which can include with locBLAST v2.0 library is below. All the programs except RPS-BLAST and RPS-TBLASTN support both online and offline sequence database search, and customized BLAST search by changing advanced algorithm parameters (Fig. 1).

**Fig 1:**
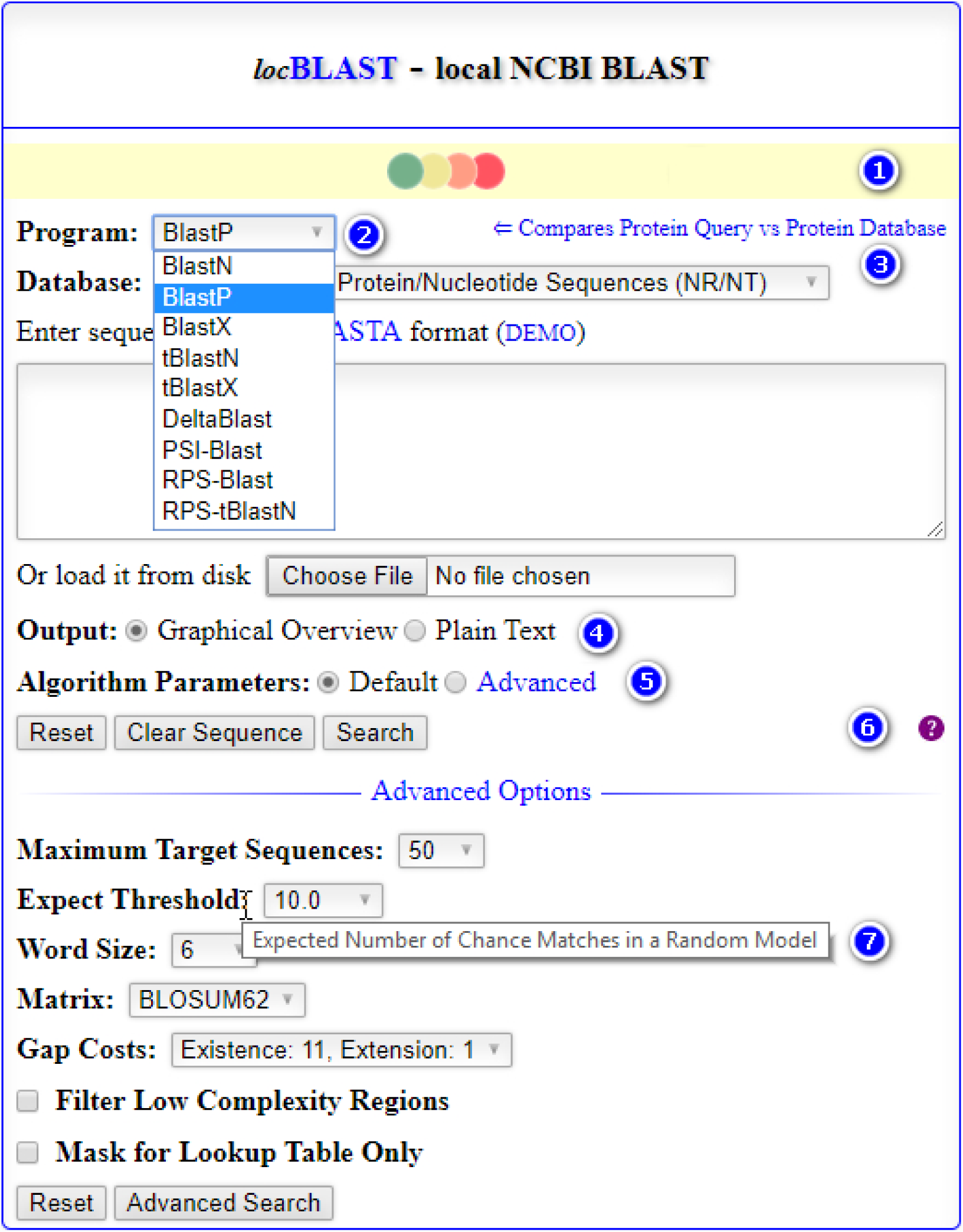
Homepage of *loc*BLAST v2.0 web form with default and advanced BLAST+ search options. (1) progress bar and status bar, (2) advanced BLAST+ programs deltablast, psiblast, rpsblast, and rpstblastn, (3) hyperlink to detailed program description page, with short title, (4) option to graphical or plain text output, (5) option to advanced search algorithm parameters, (6) tutorial page, and (7) mouse over tooltip text description of the keyword.

- BLASTN program compares the nucleotide query sequence with the nucleotide sequences in the database.
- BLASTP program compares the protein query sequence with the protein sequences in the database.
- BLASTX program translates the nucleotide query sequence and compares with the protein sequences in the database.
- TBLASTN program compares the protein query sequence with the nucleotide sequences in the database, by instantly translating the nucleotide sequences in the database before comparison.
- TBLASTX program compares the translated six reading frames of nucleotide query sequence with the translated six reading frames of nucleotide sequences in the database. The query is translated on the fly before searching sequences in the database.
- DELTA-BLAST (Boratyn et al., 2012) program compares the query protein sequence with the protein sequences in the database by constructing the position-specific scoring matrices (PSSM) from the conserved domain database search results.
- PSI-BLAST (Altschul et al., 1997) program compares the query protein sequence with the protein sequences in the conserved domain database using PSSM scores by position specific iteration method.
- RPS-BLAST program compares the query protein sequence with the protein sequences in the conserved domain database through profiles. It is a reverse of the PSI-BLAST search.
- RPS-TBLASTN program compares the translated six reading frames of nucleotide query sequence with the protein sequences in the conserved domain database through profiles.

The stand-alone NCBI BLAST+ must be installed and configured in each client machines for individual BLAST search. Moreover, the clients can’t access other databases; and specialization is needed to perform the advanced BLAST search. To outcome the issues, *loc*BLAST v2.0 is developed to host a graphical interfaced BLAST server which can accomplish online and offline database search. The default and advanced BLAST algorithm parameters in *loc*BLAST v2.0 web form automatically adjust to the default options according to the BLAST+ program and database on selection. For each BLAST+ program, it has a short title with a hyperlink to the webpage containing a detailed description of the program and algorithm parameters.

## 2. Methods

The *loc*BLAST v2.0 library is designed using PHP, CSS, and pure JavaScript. It functions under nine different phases: (i) FASTA file format validation, (ii) amino acid or nucleotide sequence validation, (iii) proper BLAST+ program parameters validation, (iv) back-end BLAST+ program execution, (v) BlastXML2 file parser (Ashok *et al*., 2017), (vi) graphical overview of hits of sequence alignments display, (vii) display tabular format short summary of hits of sequence alignments, (viii) formatted three row of sequence alignment, and (ix) save sequence to the disk. The PHP script executes the NCBI BLAST+ programs using exec() function through passing parameters from the HTML form fields. *loc*BLAST v2.0 allows users to submit input sequence either through the text area or file upload. The output of BLAST+ result is generated in plain HTML format or user interactive graphical format. In the graphical output, graphical overview (Fig. 2) and tabular summary are properly hyperlinked with tooltip text to the sequence alignments.

**Fig 2:**
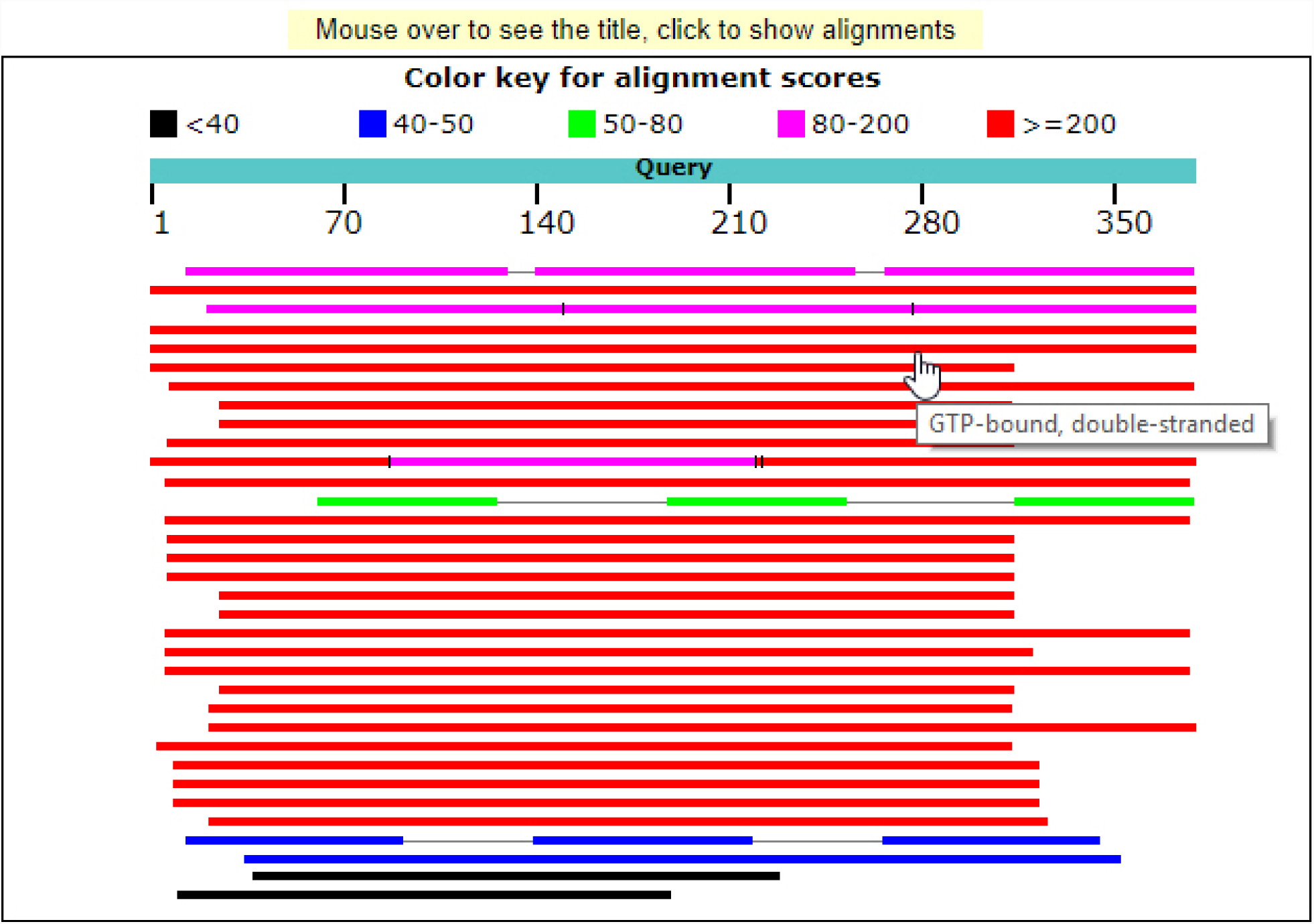
Graphical overview of hits of sequence alignment sorted by scores (color keys), similar to native NCBI BLAST result.

### 2.1 Implementation

The *loc*BLAST v2.0 library is designed user-friendly for both beginners and developers. Implementation of *loc*BLAST v2.0 to a local BLAST server consists of two simple stages: (i) Downloading and extracting files, scripts, and BLAST+ binaries from GitHub and NCBI FTP to the web directory, and (ii) Custom database preparation or modification.

#### 2.1.1 Installation and configuration

*loc*BLAST v2.0 can be easily installed in any PHP supporting webserver and platform. A brief step-wise instruction for setting *loc*BLAST v2.0 is below:

1. Create a new directory named ‘blast’ (optional) in the web-server parent directory (usually named as ‘htdocs’, ‘www’, ‘wwwroot’, or ‘webpath’).
2. Download latest BLAST+ binaries from NCBI FTP repository (ftp://ftp.ncbi.nih.gov/blast/executables/blast+/LATEST) based on the platform and extract to the ‘blast’ directory.
3. Download *loc*BLAST v2.0 from GitHub (https://github.com/AshokHub/locBLAST) and extract to the ‘blast’ directory.
4. Online biological databases can be downloaded from NCBI FTP repository (ftp://ftp.ncbi.nih.gov/blast/db) and moved to ‘db’ directory to perform database search in offline. A brief protocol for custom database preparation is explained in section 2.1.2.
5. Add or modify ‘database name’ and ‘path of the database file’ codes in ‘index.php’ and ‘footer.php’ script for the custom database accordingly (optional).
6. Open the URL http(s)://<hostname>:<port>/blast (for instance http://localhost/blast) in the web browser to enter the query sequence.
7. Choose appropriate BLAST program to use and database to search (Fig. 1).

The *loc*BLAST v2.0 provides access to search online databases namely Non-Redundant Protein/Nucleotide Sequences (NR/NT), UniProtKB/Swiss-Prot (SwissProt), Reference Proteins Sequences (RefSeq_Protein), Reference RNA Sequences (RefSeq_RNA), Expressed Sequence Tags (EST), Protein Data Bank (PDB), and Patented Protein Sequences (PAT); but not limited.

#### 2.1.2 Custom database preparation

The custom database preparation requires little technical knowledge of BLAST+ programs. BLAST+ program supports the nucleotide database, protein database, and conserved domain database search. After database preparation, the form fields to be added or modified to existing database list. A simple protocol for creating own database is below:

- For creating a nucleotide database – execute the following command-line./**makeblastdb -in** db/test_na.fas **-out** db/test_na **-dbtype nucl -title** Test_NA
- For creating a protein database – execute the following command-line./**makeblastdb -in** db/test_aa.fas **-out** db/test_aa **-dbtype prot -title** Test_AA
- For creating a RPS/PSI (CDD) database – execute the following command-line./**makeprofiledb -in** db/CDD/test_CDD_db.pn **-out** db/CDD/test_CDD_db **-dbtype rps -title** Test_CDD_db
- For creating a Delta (CDD) database – execute the following command-line./**makeprofiledb -in** db/CDD/test_Delta_db.pn **-out** db/CDD/test_Delta_db **-dbtype delta -title** Test_Delta_db

Where, the words represented in bold are keywords and should not be modified. The custom databases namely ‘Test_NA’, ‘Test_AA’, ‘Test_CDD_db’, ‘PDTDB’, and ‘Smart’ are included in the *loc*BLAST v2.0 library for testing.

#### 2.1.3 Database manipulation

The BLAST+ applications such as blastdb_aliastool, blast_formatter, blastdbcheck, convert2blastmask, dustmasker, segmasker, blastcmd, makembindex, and windowmasker are used for database manipulation or data retrieval purposes.

## 3. Results

The result page of *loc*BLAST v2.0 search produces a graphical output which contains well-formatted sequence alignment, color key, and summary table closely similar to the online BLAST result. Moreover, the web form and result page contain hyperlinks linked to the source or anchors, and keywords are annotated with a brief description of the term by tooltip text.

The graphical output consists of five section: (i) program search parameters, (ii) distribution of top BLAST hits in a graph, (iii) summary table of BLAST hits, (iv) row-wise alignment for each hits, and (v) Karlin-Altschul statistics (Fig. 3). Moreover, the database accession number and definition line in the graphical output are automatically annotated and hyperlinked to the source database entry based on the database identifiers namely *gb, emb, dbj, pir, prf, sp, pdb, pat, bbs, gnl*, and *ref*. In plain text output, the search result is presented by monospace text and interlinks to the sequence alignment hits (Fig. 4).

**Fig 3:**
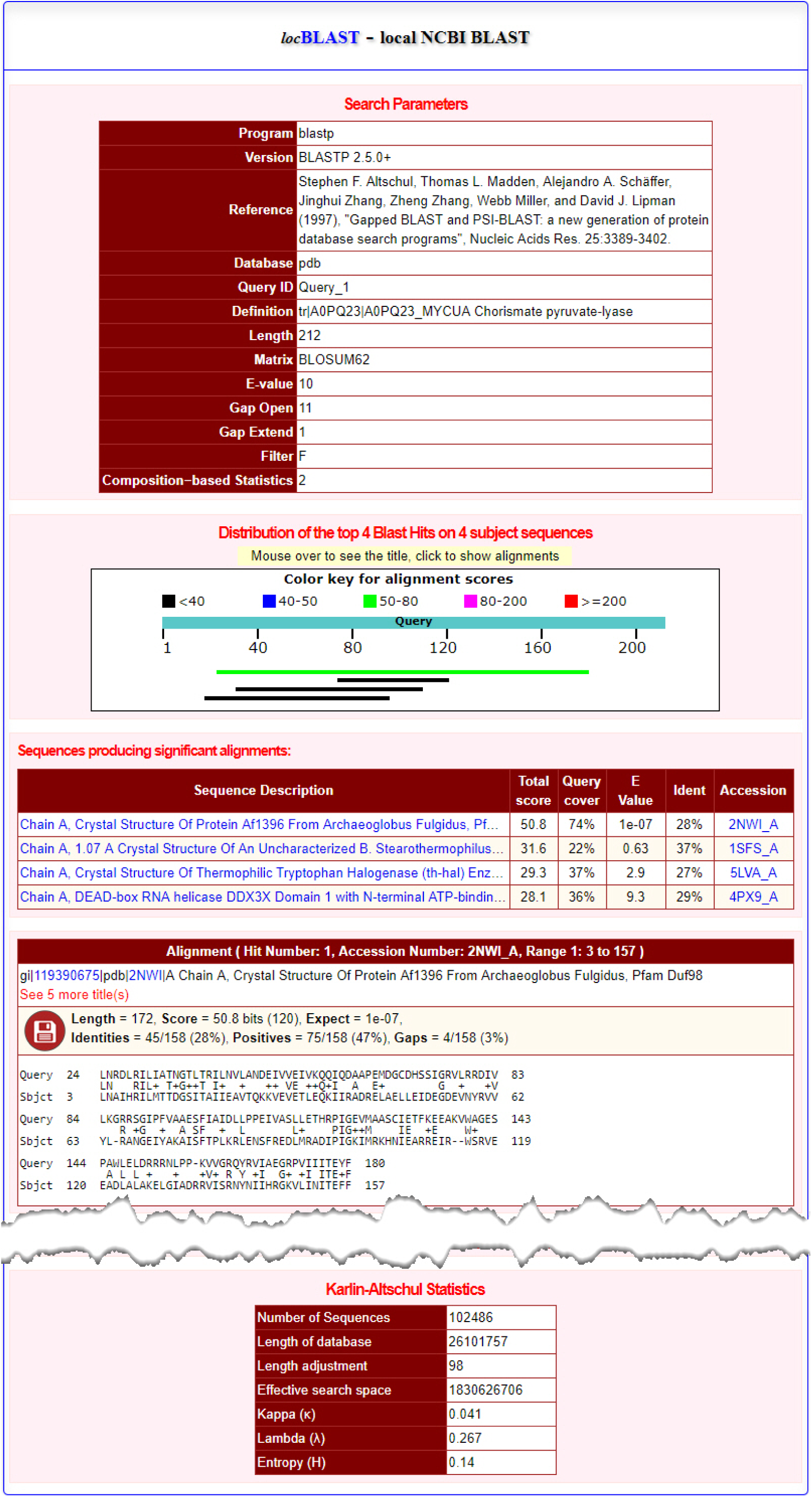
Graphical output page of *loc*BLAST v2.0 sequence search result (cropped large page).

**Fig 4:**
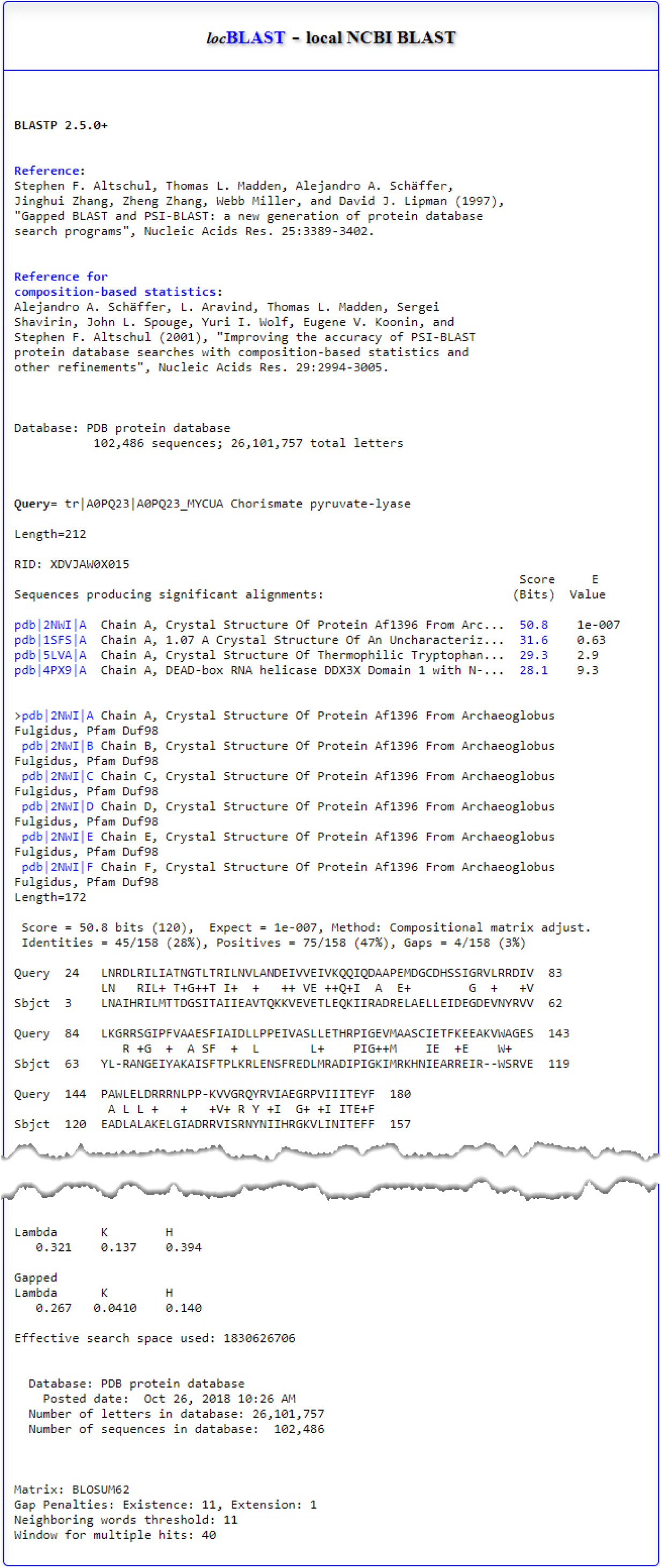
Plain text output page of *loc*BLAST v2.0 sequence search result (cropped large page).

*loc*BLAST v2.0 does not depend on any additional third-party scripts or plugins to run the NCBI BLAST+ program. It is platform independent and hence it works in most of the traditional and modern browsers. NCBI BLAST+ executables in the *loc*BLAST v2.0 can be simply updated by downloading current release from NCBI FTP site and replacing the old executables. The current stable release of NCBI BLAST+ is v2.8.1 (as on January 11, 2019), distributed as source codes and compiled binaries for Windows, Linux, and Macintosh operating systems as the installable and compressed files. The *loc*BLAST v2.0 PHP library and the test database files were freely available at GitHub under GNU General Public License v3.0.

---

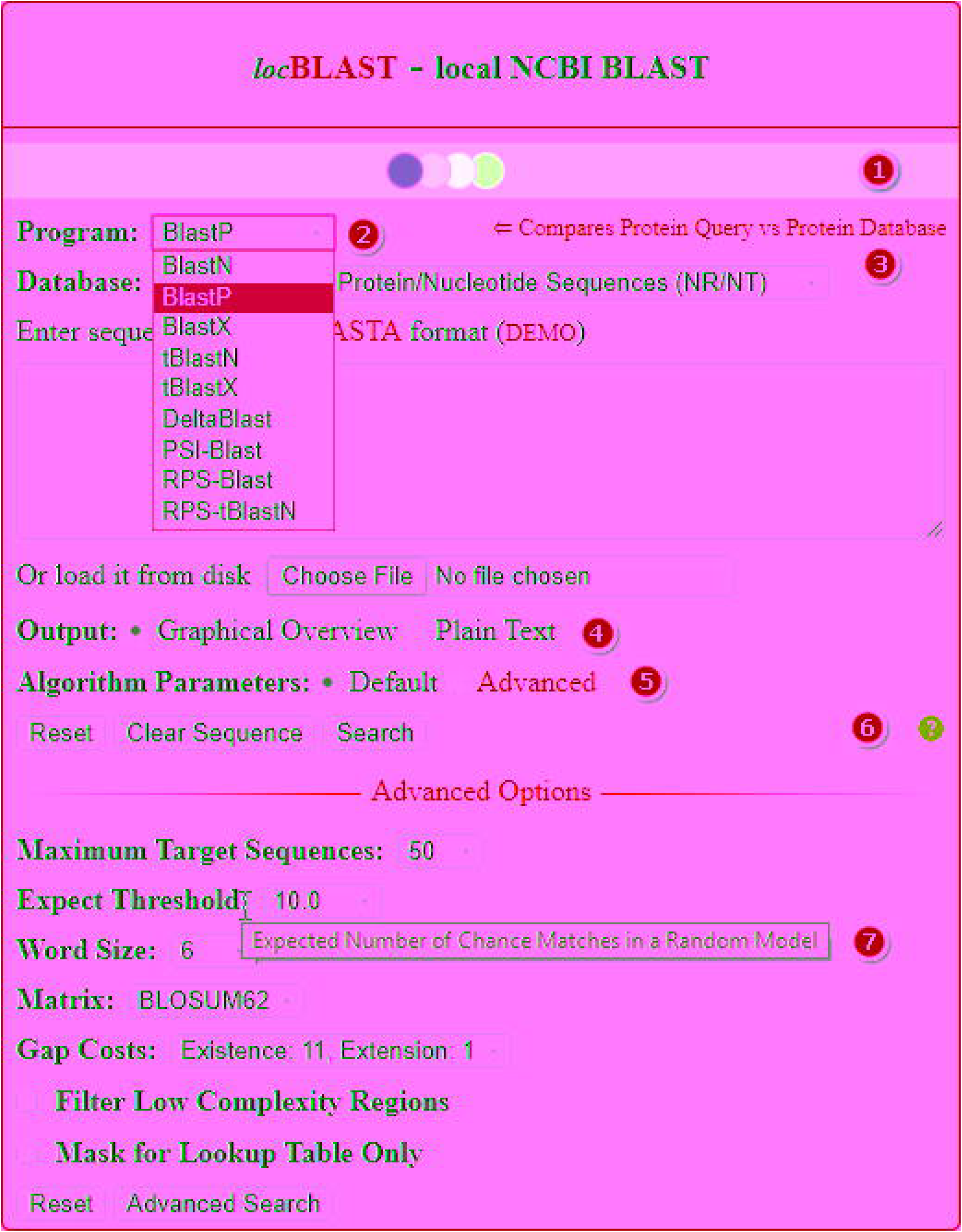

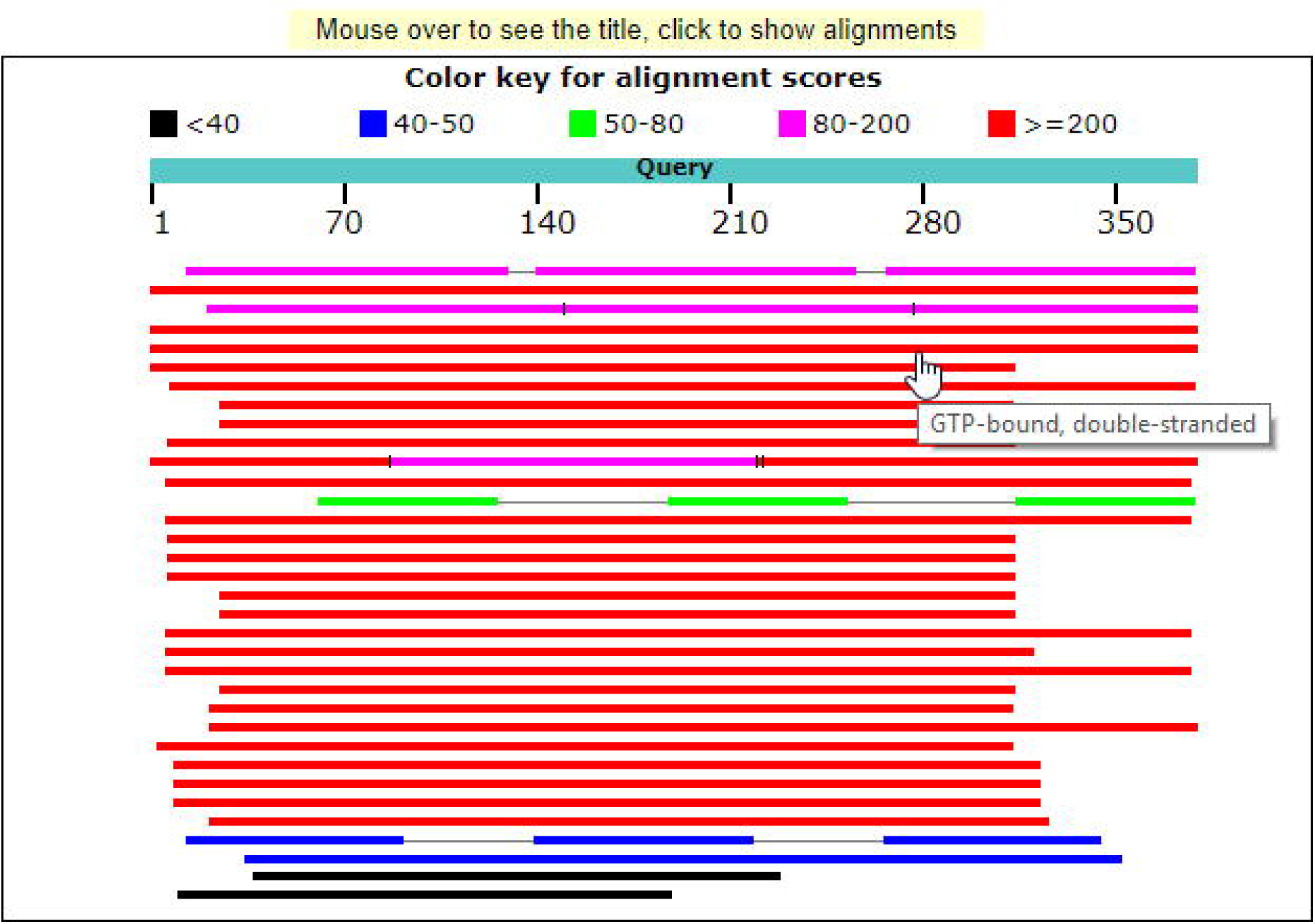

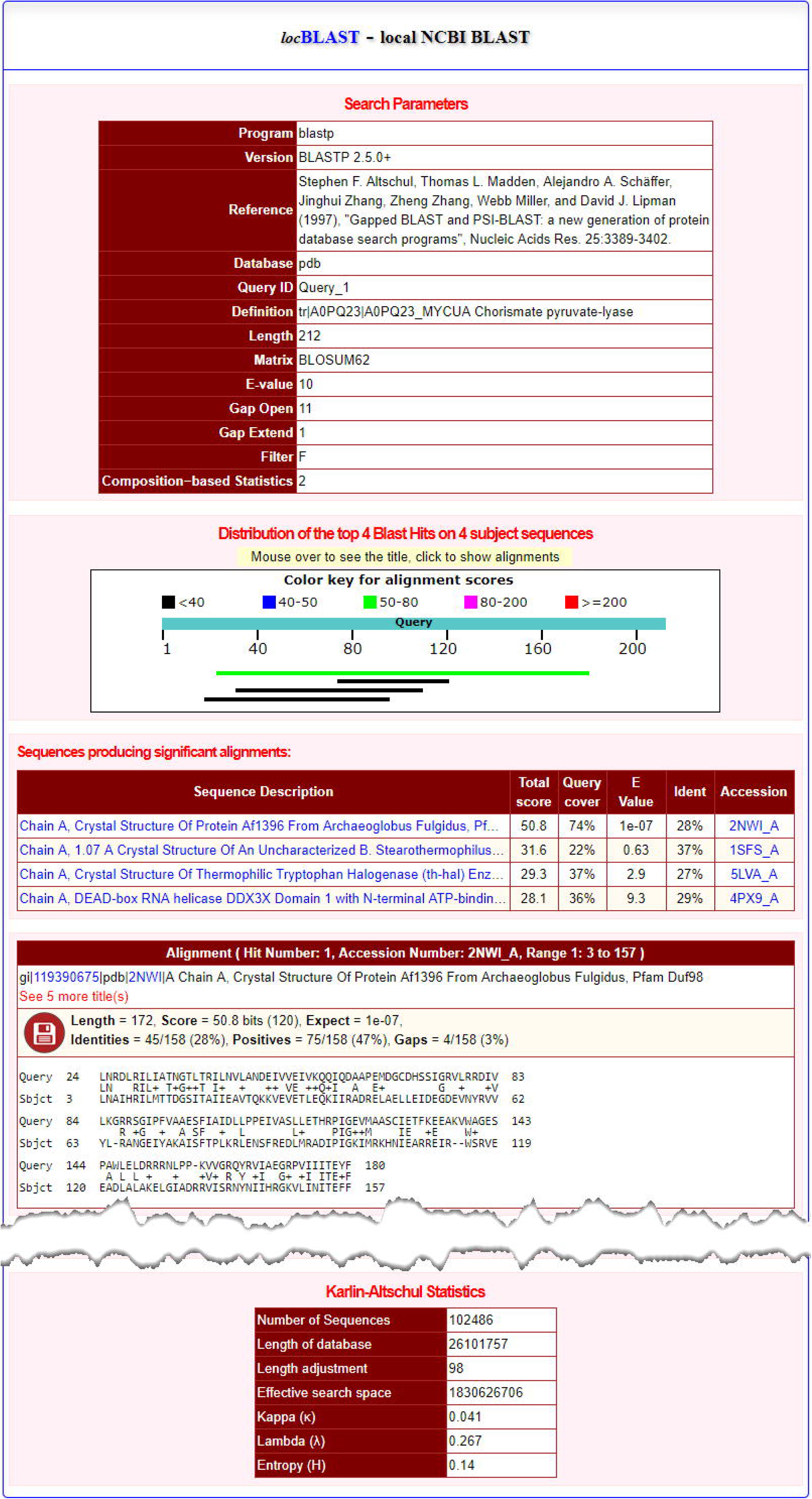

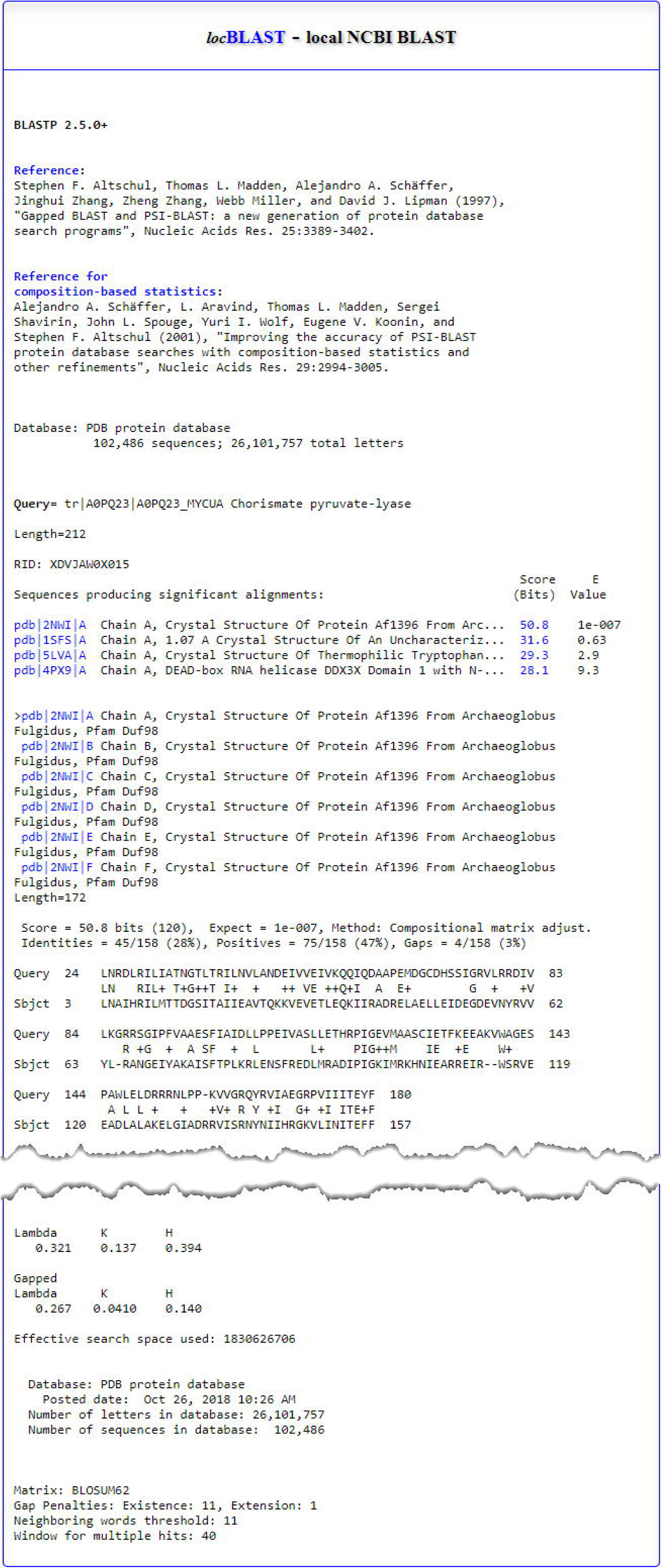

